# Synaptic proteomics identifies cathepsin B as a regulator of synapse remodelling during learning

**DOI:** 10.64898/2026.09.13.751219

**Authors:** Aelon Rahmani, Michaela E Johnson, Rosemary A Coleman, Nina Hartman, Anne Poljak, Yee Lian Chew

## Abstract

Learning requires dynamic changes in synaptic protein composition. Most synaptic proteomic studies capture endpoint snapshots following training, overlooking molecular changes occurring during learning itself. We previously established TurboID proximity proteomics as an approach to capture protein-level changes during learning in the nervous system of *Caenorhabditis elegans*. Here, we apply this strategy to synapses. Trained synapses exhibited a distinct proteome compared with mock-trained controls, with pathways enriched for neurotransmission, synaptic reorganisation, protein trafficking, and autophagy. Enrichment of autophagy proteins led us to investigate the cathepsin B orthologue CPR-4, a previously uncharacterised candidate for learning identified exclusively in trained synapses. Functional validation identified CPR-4 as a novel regulator of memory: CPR-4 is required for learning, and the absence of CPR-4 results in trained synaptic proteomes that are more like those of mock-trained synapses. Our findings support a model through which CPR-4 promotes synaptic remodelling required for learning through targeted protein clearance.

## Introduction

Learning and memory require dynamic changes in synaptic structure and function that alter communication between neurons. Behavioural plasticity is accompanied by changes in synaptic protein composition, including proteins involved in receptor trafficking, cytoskeletal organisation, neurotransmission, and local protein synthesis [9–12]. A well-characterised example is the increased expression and synaptic localisation of AMPA-type glutamate receptors during long-term potentiation [13–15]. While many studies have identified individual proteins required for learning, synaptic plasticity is ultimately mediated by coordinated changes across protein networks. Defining how these networks are remodelled during learning remains a central challenge in understanding memory formation.

Proteomic approaches have provided important insights into the molecular composition of synapses and how they change with learning. Most studies combine synaptic fractionation methods, such as synaptosome or postsynaptic density enrichment, with mass spectrometry to identify proteins associated with learning-related plasticity [11, 12, 17]. However, because they typically profile tissue collected after training has occurred, they provide only a snapshot of the synaptic proteome at a single timepoint. Given that many proteins dynamically traffic between subcellular compartments during memory formation, approaches capable of capturing proteins within their native cellular context during learning may reveal additional mechanisms underlying synaptic plasticity [19].

TurboID proximity labelling provides a powerful strategy as it can define local protein networks *in vivo* within a desired time frame. TurboID is an engineered biotin ligase that rapidly biotinylates proteins within approximately 10 nm, enabling their subsequent identification by mass spectrometry [20]. By targeting TurboID to specific tissues or subcellular compartments, proteomes can be analysed with both spatial and temporal precision, making it particularly well suited for studying dynamic biological processes in genetically tractable organisms such as *C. elegans* [1, 6, 21, 22]. We recently used this approach to identify proteins present in *C. elegans* neurons during associative learning, generating a dynamic proteomic profile of memory encoding [23].

Here, we use synapse-targeted proximity proteomics to define the protein networks present at synapses during memory encoding. We identified a learning-associated synaptic proteome comprising >300 proteins exclusively detected during memory encoding. This proteome was enriched for pathways involved in protein expression, trafficking, and degradation, highlighting proteostasis as a prominent feature of synaptic remodelling in learning. Functional analyses identified the cathepsin B (CTSB) orthologue CPR-4 as a previously unrecognised regulator of gustatory learning. We further show that CPR-4 is required for changes in synaptic protein composition due to learning, including regulation of the active-zone protein ELKS-1. Together, our findings support a model in which memory formation requires active remodelling of the synaptic proteome through proteostasis mechanisms, including protein clearance.

## Results

### Synaptic content changes during appetitive learning in *C. elegans*

To define the protein network present in *C. elegans* synapses during memory formation, we took advantage of the spatiotemporal controllability of TurboID proximity labelling [23]. We targeted TurboID to the presynaptic active zone through its fusion with ELKS-1, a scaffolding protein required for synapse organisation [1, 24, 25]. We subjected ELKS-1::TurboID transgenic animals to a gustatory associative learning paradigm [23], where the TurboID substrate (biotin) was provided only during training and depleted at all other time points, including during development (see *Methods*, **Fig. 1A**). This strategy enabled temporal control of proximity labelling during memory encoding, generating what we refer to as the learning proteome [23].

**Figure 1.**
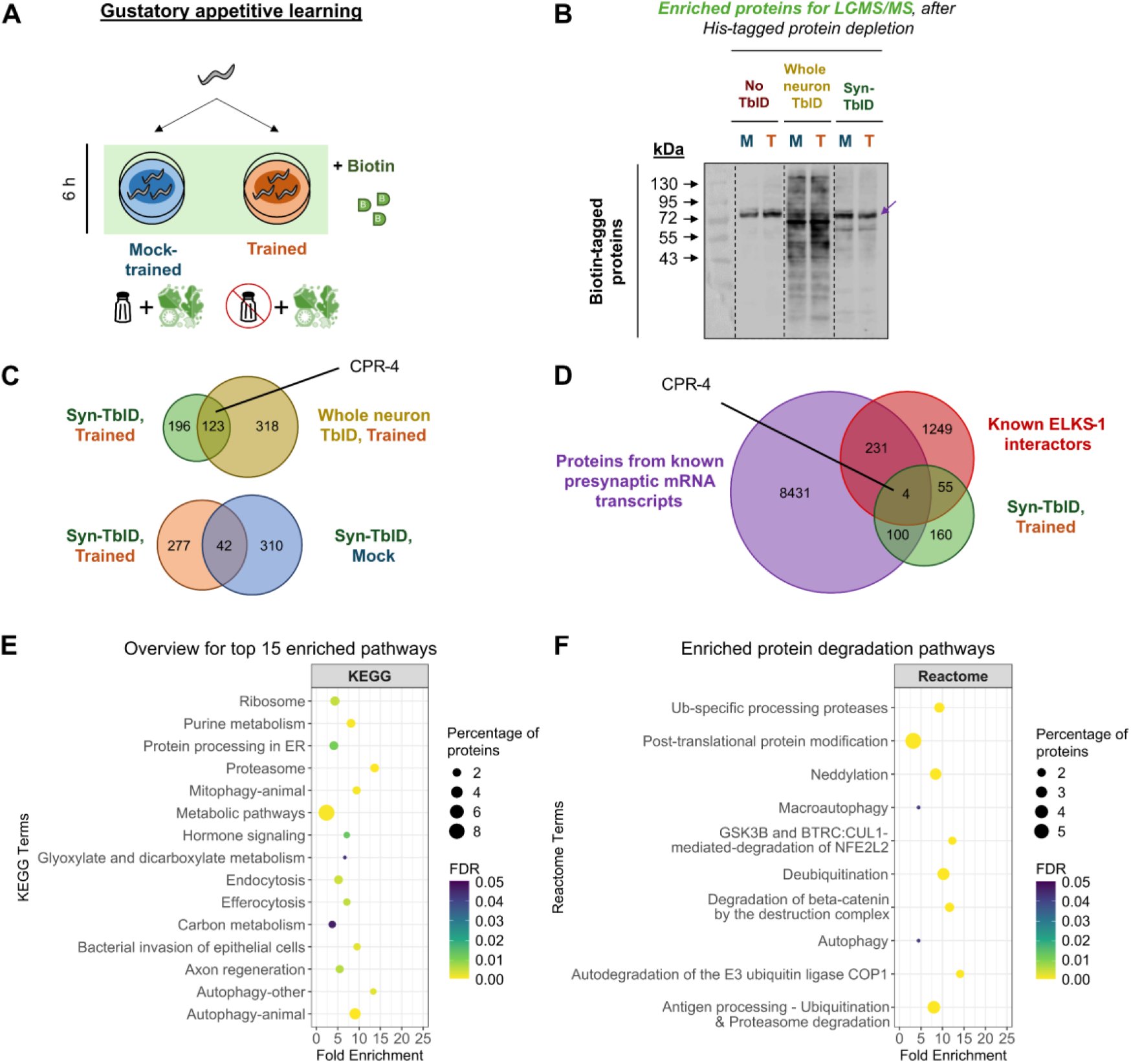
Appetitive gustatory learning changes proteins and molecular pathways present at synapses. **(A)** Schematic showing our biotin treatment strategy. Biotin (in green) was supplied only during the 6 h mock-training (salt + food) or training (no salt + food) period. **(B)** Western blots of biotinylated proteins enriched by double pull-down for LC−MS/MS. M = mock-trained; T = trained; TbID = TurboID; Syn-TbID = synaptic TurboID. **(C)** Comparison of proteins detected in different TurboID proteomes. Percentage overlap indicates the proportion of proteins shared between two proteomes relative to the total proteins detected across both datasets. CPR-4, explored in detail in this study, is highlighted here. **(D)** Comparison of the synaptic learning proteome with published datasets of presynaptically localised transcripts and known or predicted ELKS-1 interactors. CPR-4 was one of the proteins shared between all three datasets and was selected for further investigation. **(E, F)** Pathway enrichment analyses of the synaptic learning proteome. Fold enrichment was calculated relative to all annotated *C. elegans* proteins, and percentage of proteins refers to the proportion of the 319 proteins from trained synapses associated with each pathway. False discovery rate (FDR) ≤ 0.05. Plots were created using R (v4.6). **(E)** KEGG pathways ranked by percentage of proteins. **(F)** Selected Reactome pathways associated with protein degradation and proteostasis.

To identify proteins associated with synapses during learning, we compared our line with ELKS-1-tagged TurboID (‘synaptic TurboID’) with two control lines: (1) animals not expressing TurboID (‘no TurboID’), and (2) animals expressing untagged TurboID, i.e. not targeted to a specific subcellular region (‘whole neuron TurboID’). All three lines showed intact gustatory appetitive learning under restricted biotin conditions: with trained animals displaying reduced attraction to high salt compared with naïve and mock-trained controls (**Fig. S1**). Following the training (or mock-training) protocol, biotinylated proteins were enriched and analysed by mass spectrometry (see *Methods*). Consistent with successful TurboID-dependent biotinylation, we observed a stronger biotinylation signal from TurboID-expressing samples compared with no TurboID groups (**Figs. 1B & S2**).

After subtracting proteins detected in the corresponding no-TurboID controls (*n = 3*), four proteomes were generated: whole-neuron mock-trained (269 proteins), whole-neuron trained (442 proteins), synaptic mock-trained (352 proteins), and synaptic trained (319 proteins; hereafter referred to as the synaptic learning proteome) (**Table S2**). Although substantial overlap was observed between the trained whole-neuron and synaptic proteomes (123 proteins; ∼19%; **Fig. 1C**), likely reflecting the inclusion of synaptic proteins within the broader neuronal proteome, synaptic TurboID additionally identified 196 proteins that were not detected using whole-neuron TurboID (**Fig. 1C**). Separately, comparison of trained and mock-trained synaptic proteomes revealed only 42 shared proteins (∼7%, **Fig. 1C**). This indicates that synapses undergo extensive remodelling in response to learning, consistent with [11, 12, 17]. Together, these findings indicate that memory encoding is accompanied by substantial changes in the protein network present at synapses.

We next assessed whether the synaptic learning proteome was enriched for neuronal and synaptic proteins. Analysis using the CeNGEN single neuron transcriptomics database [26], showed strong enrichment for neuronal expression, with ≥94% of proteins in each TurboID dataset expressed in the nervous system [27] [accessed 11/07/2026]. Consistent with successful synaptic targeting during learning, Gene Ontology analysis (Biological Process) identified enrichment for neurotransmission, synaptic vesicle function, and synapse organisation pathways (**Fig. S3**) [28, 29]. In addition, in trained synapses, 104 proteins (∼33%) correspond to mRNA transcripts previously reported to localise to presynapses [30], while 59 (18%) proteins overlapped with known ELKS-1 interactors [1, 24, 31–35] (**Fig. 1D**). Consistent with localisation to the ELKS-1-containing presynaptic active zone, trained synapses also contained seven PDZ domain proteins (∼11% of all PDZ domain proteins encoded in the worm genome) (**Table S3**), including predicted ELKS-1 interactors CNK-1/CNKSR2/3 and PTP-1/PTPN4 [36, 37], which are homologs of mammalian learning regulators [38, 39]. Together, these analyses support the successful capture of proteins associated with synapses during learning.

We next identified the most highly-represented molecular pathways in the synaptic learning proteome by KEGG pathway analysis [40, 41]. We observed 19 enriched terms (**Table S4**), which can be broadly categorised into three functional categories: (1) protein expression and trafficking (2) cell signalling regulation and (3) protein degradation pathways (**Fig. 1E**). Notably, several proteostasis-related pathways were enriched, including autophagy, proteasomal degradation, and efferocytosis. Similar results were obtained using the Reactome pathway database [42] (**Table S5**), which also identified autophagy and proteasome-related processes. These findings indicate that gustatory appetitive learning is accompanied by coordinated changes in synaptic proteins associated with protein trafficking, turnover, and degradation.

To move beyond pathway-level analyses and to identify specific proteins that may contribute to learning-induced synaptic changes, we examined proteins uniquely detected in trained synapses. Based on KEGG pathway analysis, we focussed on endocytosis (7 proteins, **Fig. 2A**), purine metabolism (7 proteins, **Fig. 2B**), and autophagy (15 proteins, **Fig. 2C**). Many of these proteins have established roles in synaptic function and behavioural plasticity. The endocytosis network included RHO-1/Rho GTPase, which regulates cholinergic signalling in presynapses [43], and proteins implicated in synaptic vesicle recycling, dynamin DYN-1 [44] and adaptin DPY-23 [45] (**Fig. 2A**). The purine metabolism network contains guanylyl cyclases required for gustatory plasticity (GCY-33) [46, 47] and behavioural choice between conflicting stimuli (GCY-28, [7]) (**Fig. 2B**). Notably, the autophagy-associated network contained three proteins implicated in specifying cargo uptake for autophagosomes (PEK-1/PERK, ATG-16.1/ATG16, & ATG-16.2/ATG16) [40, 41, 48, 49], and three lysosomal protease enzymes (ASP-3/CTSD, CPR-1/CTSB, & CPR-4/CTSB) (**Fig. 2C**) [50, 51].

**Figure 2.**
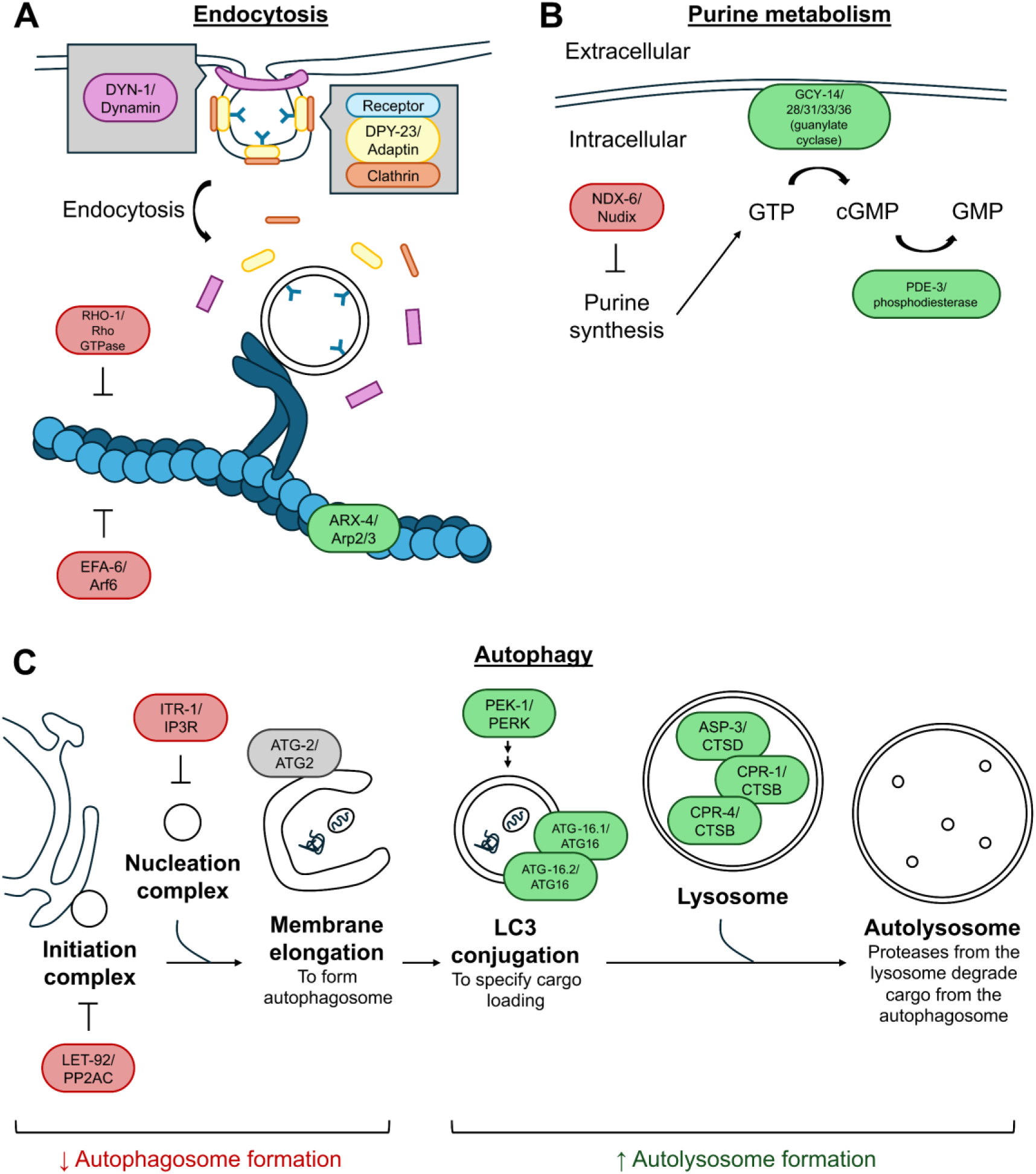
Endocytosis, purine metabolism, and autophagy as enriched within the synaptic learning proteome. **(A)** Endocytosis-associated proteins implicated in endocytic vesicle formation, trafficking, cytoskeletal regulation, and receptor internalisation are highlighted. Proteins that restrict cytoskeleton growth are in red [2], and those that promote its construction are in green [4]. **(B)** Purine metabolism-related proteins that restrict purine metabolism are in red [5, 6] and those that promote it are in green [7, 8]. **(C)** Autophagy-associated proteins: Negative regulators of autophagosome formation are in red, whereas proteins associated with autolysosome formation are in green. This schematic is adapted from [18], and KEGG pathway diagrams on ShinyGO (v0.85). Schematics show proteins detected exclusively in trained synapses of *C. elegans*. Each node represents a ‘*C. elegans* protein name’/‘Human homolog name’.

Autophagy is mainly known for its role in degrading damaged proteins and organelles, and recovering their components for reuse (reviewed in [52]). However, it is increasingly recognised for its role in presynapse organisation [53–55], synaptic proteome content modulation [56, 57], synapse morphology [58], and learning [18, 56, 58, 59]. The enrichment of autophagy-associated proteins within trained synapses therefore suggests that protein clearance pathways may contribute to learning-dependent remodelling of the synaptic proteome. Importantly, many of the autophagic components detected have not previously been implicated in learning. We therefore considered the synaptic learning proteome to be a potential source of novel regulators connecting proteostasis to memory formation and selected one such candidate, CPR-4, for further investigation.

### Gustatory learning in the worm is dependent on cathepsin B ortholog CPR-4

The enrichment of autophagy-related proteins in trained synapses prompted us to investigate the functional significance of CPR-4, a CTSB orthologue. In *C. elegans,* although CPR-4 has mainly been studied for its role as a stress signal due to UV exposure or overcrowding [3, 60], several observations suggest it may also function in neuronal plasticity. First, we detected CPR-4 specifically in trained proteome datasets for synapses and whole neurons (**Figs. 1 & 2**) [23]. Second, transcriptomic studies highlight that *cpr-4* transcripts are expressed in neurons and localised to the presynapse [27, 30] (**Fig. 1D**). Finally, CPR-4 contains conserved catalytic residues present in mammalian CTSB that are required for protein degradation within autophagic pathways [3], a process increasingly recognised as an important regulator of synaptic protein turnover [56, 57]. Together, these observations suggest that CPR-4 may represent a previously unrecognised regulator of learning-dependent synaptic remodelling.

To determine whether CPR-4 is required for gustatory learning, we first examined two putative loss-of-function *cpr-4* alleles, *tm3718* (406 bp deletion and 12 bp insertion) and *ok3413* (∼ 300 bp deletion, unmapped), each of which remove roughly 30% of the gene’s single exon [61, 62]. Neither mutation affected baseline salt chemotaxis, as naïve and mock-trained mutants exhibited high-salt attraction comparable to wild-type (WT) animals (**Fig. 3A,B**). This indicates that *cpr-4* mutants do not show gross locomotor or sensory defects that could confound interpretation of learning phenotypes. Successful learning in this assay is reflected by a shift in chemotactic behaviour between mock-trained and trained animals, as quantified by chemotaxis indices (CIs). This allows learning performance to be compared between mutant and wild-type (WT) strains.

**Figure 3.**
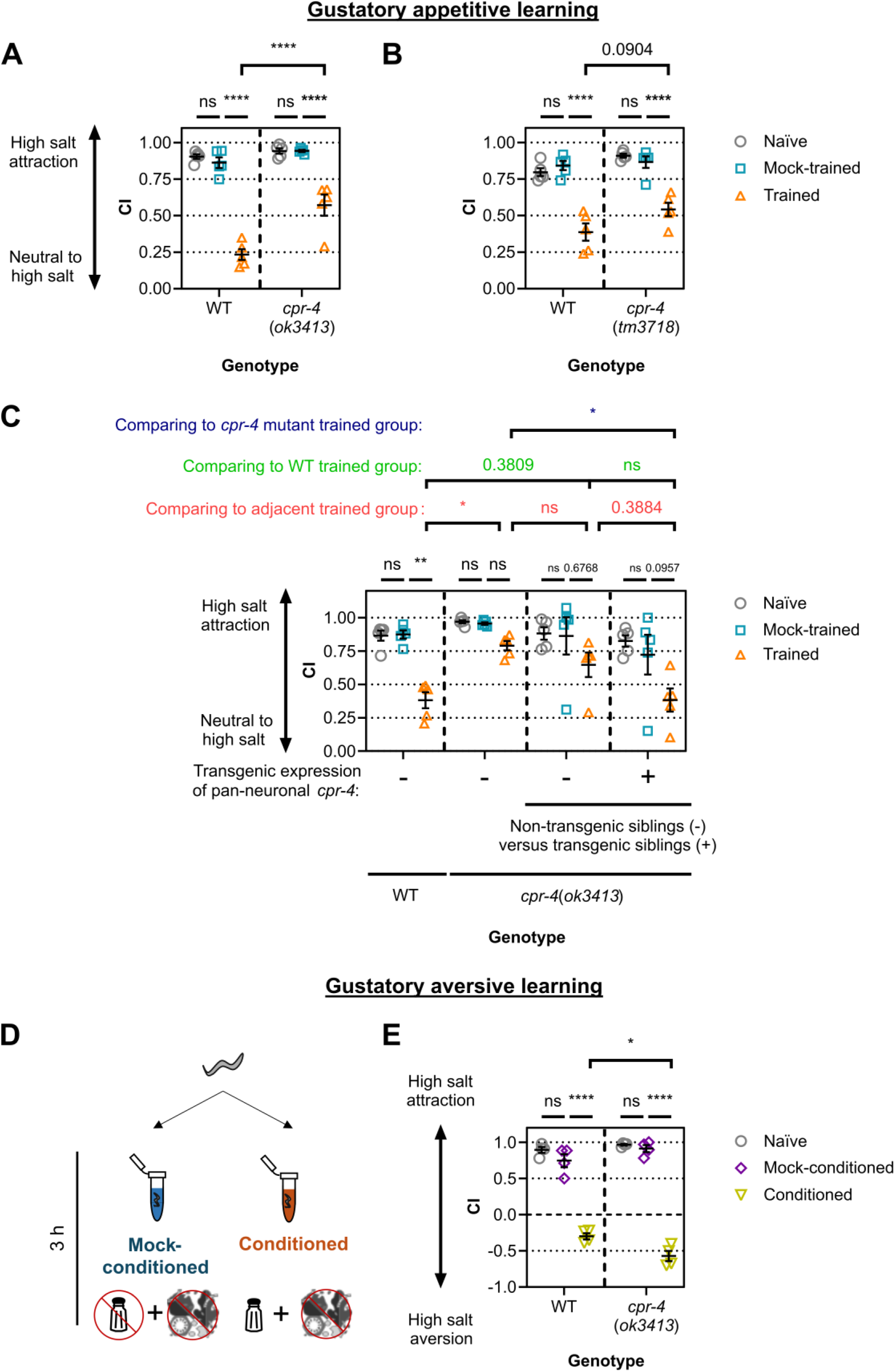
CPR-4 regulates both appetitive and aversive gustatory learning. **(A)** Chemotaxis indices for the *cpr-4(tm3718)* allele in gustatory appetitive learning. WT animals (line N2) were compared with *cpr-4(tm3718)* mutants (line CUI1911). **B)** Chemotaxis indices for the *cpr-4(ok3413)* allele in gustatory appetitive learning. WT animals (line N2) were compared with *cpr-4(ok3413)* mutants (line RB2473). **(C)** Chemotaxis indices following neuronal CPR-4 re-expression in gustatory appetitive learning. WT, *cpr-4(ok3413)* mutants, and animals carrying the pan-neuronal CPR-4 rescue transgene were analysed. Non-transgenic siblings are indicated by "−" and transgenic siblings by "+". The extrachromosomal array re-expresses CPR-4 pan-neuronally in transgenic animals only. Across biological replicates, 26-41% of animals were transgenic. **(D)** Schematic for gustatory aversive learning. Animals were mock-conditioned (no salt + no food) or conditioned (salt + no food). **(E)** Chemotaxis indices for the *cpr-4(ok3413)* allele in gustatory aversive learning. WT animals (line N2) were compared with *cpr-4(ok3413)* mutants (line RB2473). Error bars = mean ± SEM. Each data point represents one biological replicate (*n*). For **(A-C)**, *n* = 5 biological replicates with 2-3 technical replicates per biological replicate; some technical replicates were excluded due to insufficient sample size (<20 worms). *n* = 4 for **(E)** with 3 technical replicates per biological replicate. Each technical replicate contained 22-314 worms. Statistical analyses: Two-way ANOVA with Tukey’s multiple comparisons test (**** ≤ 0.0001, * ≤ 0.05, ns = non-significant; all other p-values shown).

Appetitive gustatory learning was impaired in *ok3413* mutants, as trained *ok3413* animals exhibited significantly higher chemotaxis indices (CI) than trained WT controls (**Fig. 3B**). Although a similar effect was observed in *tm3718* mutants, this did not reach statistical significance (**Fig. 3A**). As the *ok3413* allele produced the stronger learning phenotype, we therefore continued testing only *ok3413* animals in subsequent assays.

To determine whether CPR-4 acts within the nervous system, we re-expressed CPR-4 pan-neuronally in the *cpr-4(ok3413)* background. Neuronal re-expression of CPR-4 significantly rescued the learning defect observed in *cpr-4(ok3413)* mutants, restoring performance to WT levels (**Fig. 3C**). This was indicated by two comparisons: (1) trained transgenics showed significantly lower CIs than mutants and (2) trained transgenic animals performed indistinguishably from trained wild-type controls (**Fig. 3C**). Together, these findings identify CPR-4 as a neuronal regulator of appetitive gustatory learning.

Many proteins exert different effects on learning depending on valence [63]. We next assessed *ok3413* animals in an aversive gustatory learning paradigm, where animals learn to avoid high salt in response to paired exposure to salt and food deprivation (**Fig 3D**) [64]. Interestingly, the learnt aversive response in this protocol was enhanced in *ok3413* mutants compared with WT controls, with mutants showing a greater CI change after learning (**Fig. 3E**). Together, these data show that CPR-4 influences gustatory learning in a valence-dependent manner, promoting appetitive learning while limiting aversive learning.

### CPR-4 is required for learning-dependent remodelling of the synaptic proteome

CPR-4 is predicted to act as a lysosomal protease [3], raising the possibility that learning defects in *cpr-4* mutants may arise from impaired clearance of synaptic proteins during learning. We therefore hypothesised that CPR-4 contributes to learning-dependent remodelling of the synaptic proteome. In this model, proteins present at synapses under basal conditions may need to be selectively degraded. For example, proteomic analysis of aversive learning in mice found that more synaptic proteins were downregulated than upregulated following training [17]. If CPR-4 promotes learning-associated synaptic remodelling, then trained *cpr-4* mutants should fail to acquire the normal synaptic learning proteome (**Fig. 4A**). To test this prediction, we used TurboID proximity labelling to identify learning-associated changes in the synaptic protein network in the presence or absence of CPR-4.

**Figure 4.**
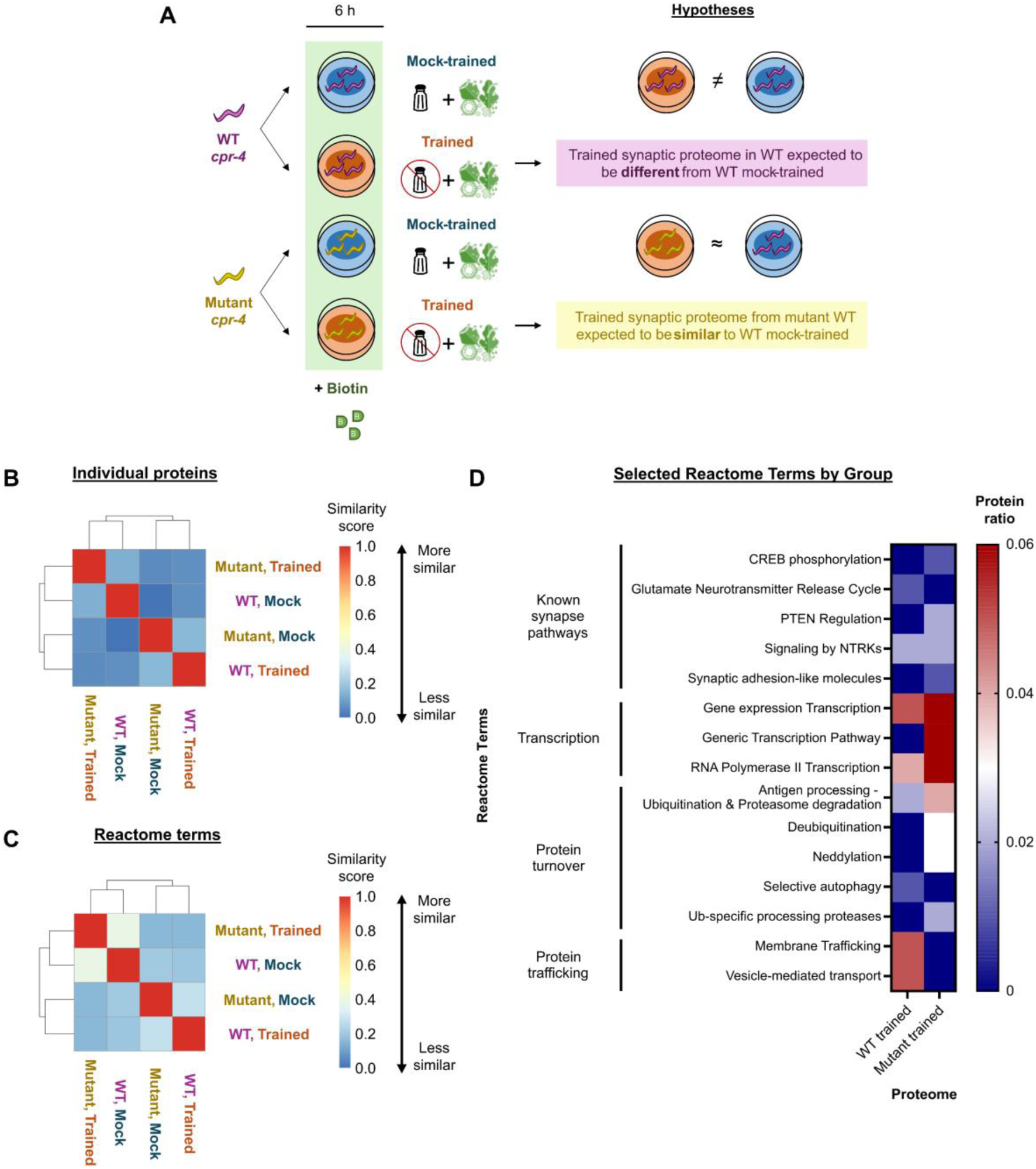
CPR-4 is required for learning-dependent remodelling of the synaptic proteome. **(A)** Experimental design and hypotheses. Synaptic TurboID was used to compare WT and *cpr-4(ok3413)* animals following mock-training or appetitive gustatory learning. For each genotype, biotin was restricted to the 6 h-period of mock-training or training. Based on its predicted lysosomal protease activity and sequence similarity to CTSB [3], we hypothesised that loss of CPR-4 would impair learning-associated remodelling of the synaptic proteome. **(B, C)** Jaccard similarity analyses comparing proteomes across genotypes and training conditions. Similarity was assessed using **(B)** proteins detected by synaptic TurboID and **(C)** Reactome pathways enriched from those proteins. Jaccard similarity coefficients were calculated using R (v4.6). **(D)** Comparison of selected Reactome pathways enriched in WT and *cpr-4* mutant learning proteomes. Enriched pathways (FDR ≤ 0.05) were exported from ShinyGO (v0.85) and visualised with GraphPad Prism (v8.0.2). Protein ratio indicates the proportion of proteins within each learning proteome associated with the indicated pathway.

We crossed the *cpr-4(ok3413)* allele into ELKS-1::TurboID transgenic worms and used proximity proteomics to compare four experimental groups. The groups differed by *cpr-4* genotype (WT or *ok3413*) and training condition (mock-trained or trained) (**Table S6**). As with previous experiments, animals were given biotin only during training (or mock-training), restricting TurboID labelling to proteins in synapses during this window (**Figs. 4A**). Consistent with previous experiments, trained *cpr-4(ok3413)* animals displayed significantly weaker appetitive learning than trained WT controls (**Fig. S4A**). All groups showed robust biotinylation after enrichment, confirming successful TurboID labelling (**Figs. S4B**). We therefore proceeded with mass spectrometry analysis.

We first tested the prediction that trained *cpr-4* mutants would fail to acquire the WT learning proteome. Consistent with this prediction, WT and *cpr-4* mutant learning proteomes shared only 27 proteins (5.5% overlap) (**Fig. S4C**, left panel). In contrast, the *cpr-4* mutant learning proteome displayed greater similarity to the WT mock-trained proteome, sharing 75 proteins (13.6% overlap) (**Fig. S4C**, right panel). Hierarchical clustering based on Jaccard similarity coefficients revealed that trained *cpr-4* mutants clustered more closely with WT mock-trained animals than with WT trained animals (**Fig. 4B**). Interestingly, WT and *cpr-4* mock-trained proteomes also differed from one another, suggesting that CPR-4 influences synaptic protein composition even in the absence of training (**Fig. 4B**). Nevertheless, the proteome of trained *cpr-4* mutants more closely resembled WT mock-trained animals than WT trained animals, suggesting that loss of CPR-4 prevents normal learning-associated proteome remodelling and promotes retention of a pre-learning synaptic state.

To determine whether these relationships extended beyond individual proteins to broader biological functions, we compared Reactome pathway enrichments across the four proteomes. Like the single protein-level analysis, the *cpr-4* trained proteome showed the greatest pathway-level similarity to the WT mock-trained proteome, sharing 115 Reactome terms (44.2%) (**Table S7**). By comparison, WT mock-trained and WT trained proteomes shared 62 terms (24.2%, **Tables S7 & S8**), mutant mock-trained and mutant trained proteomes shared 49 terms (20.1%, **Table S9**), and WT and mutant learning proteomes shared only 33 terms (17.0%, **Table S8**). Hierarchical clustering of pathway-level Jaccard similarity coefficients again grouped trained *cpr-4* mutants with WT mock-trained animals rather than WT trained animals (**Fig. 4C**). Thus, both protein- and pathway-level analyses support the conclusion that loss of CPR-4 impairs the transition to a learning-associated synaptic state.

To identify biological processes associated with CPR-4-dependent synaptic remodelling, we next examined pathways uniquely enriched in WT and mutant learning proteomes after removing the 27 proteins shared between groups (**Fig. S4C**, left panel). Reactome enrichment identified 72 pathways in the WT learning proteome and 123 pathways in the *cpr-4* mutant learning proteome (**Table S8**). To identify pathways potentially associated with CPR-4-dependent synaptic remodelling, we calculated the proportion of proteins assigned to each Reactome pathway for each learning proteome. Pathways showing differences between WT and *cpr-4* mutant learning proteomes were selected for visualisation in **Figure 4D**. Several transcription-related pathways, including Gene Expression Transcription, Generic Transcription Pathway, and RNA Polymerase II Transcription, were relatively overrepresented in the mutant learning proteome (**Fig. 4D**). Proteins contributing to these enrichments included ABL-1 and SPT-5, both previously implicated in memory-associated transcriptional responses [65]. Pathways associated with protein turnover and protein trafficking were also differentially represented between WT and mutant learning proteomes (**Fig. 4D**). For example, representation of proteasome-related pathways increased while autophagy-associated terms decreased in trained synapses in the absence of CPR-4 (**Fig. 4D**). Because proteasomal degradation, autophagy, and cathepsin B activity are functionally linked in *C. elegans* [66], these changes suggest that loss of CPR-4 alters protein turnover processes associated with learning-dependent synaptic remodelling. Together, these findings identify transcriptional regulation, protein turnover, and protein trafficking as candidate processes through which CPR-4 promotes learning-dependent remodelling of the synaptic proteome.

### Learning-associated reduction of synaptic ELKS-1 requires CPR-4

To validate our proteomic observations at the level of an individual synaptic protein, we quantified fluorescence intensity of mNeonGreen tagged ELKS-1 by fluorescence microscopy. Learning relies on protein trafficking to synapses [19], and pathway analyses identified protein trafficking as a candidate process affected by loss of CPR-4 (**Fig. 4D**). Furthermore, transcription of the mammalian ELKS-1 ortholog (*Erc2*) has been reported to increase following aversive learning in mice [67]. We therefore examined whether ELKS-1 abundance changes during appetitive gustatory learning and whether any such changes depend on CPR-4 to further build our model mechanism.

We first quantified ELKS-1 fluorescence in the *C. elegans* nerve ring (**Fig. 5A**), a major synaptic hub, in WT animals subjected to mock-training or appetitive gustatory learning (**Figs. 5B**). Surprisingly, trained WT animals showed significantly lower ELKS-1 fluorescence levels than mock-trained controls (**Figs. 5C & 5D**), indicating that learning is associated with a reduction in synaptic ELKS-1 abundance. Given that CPR-4 is enriched within the ELKS-1 proximal proteome during learning (**Fig. 2C**) and is required for learning-associated synaptic remodelling (**Fig. 4**), we next tested whether this decrease depended on CPR-4 function. Consistent with this possibility, trained *cpr-4(ok3413*) animals failed to exhibit the learning-associated decrease in ELKS-1. Instead, ELKS-1 levels in trained mutants remained comparable to those of mock-trained animals (**Fig. 5C & 5D**). These findings identify ELKS-1 as a candidate target of CPR-4-dependent synaptic remodelling during learning.

**Figure 5.**
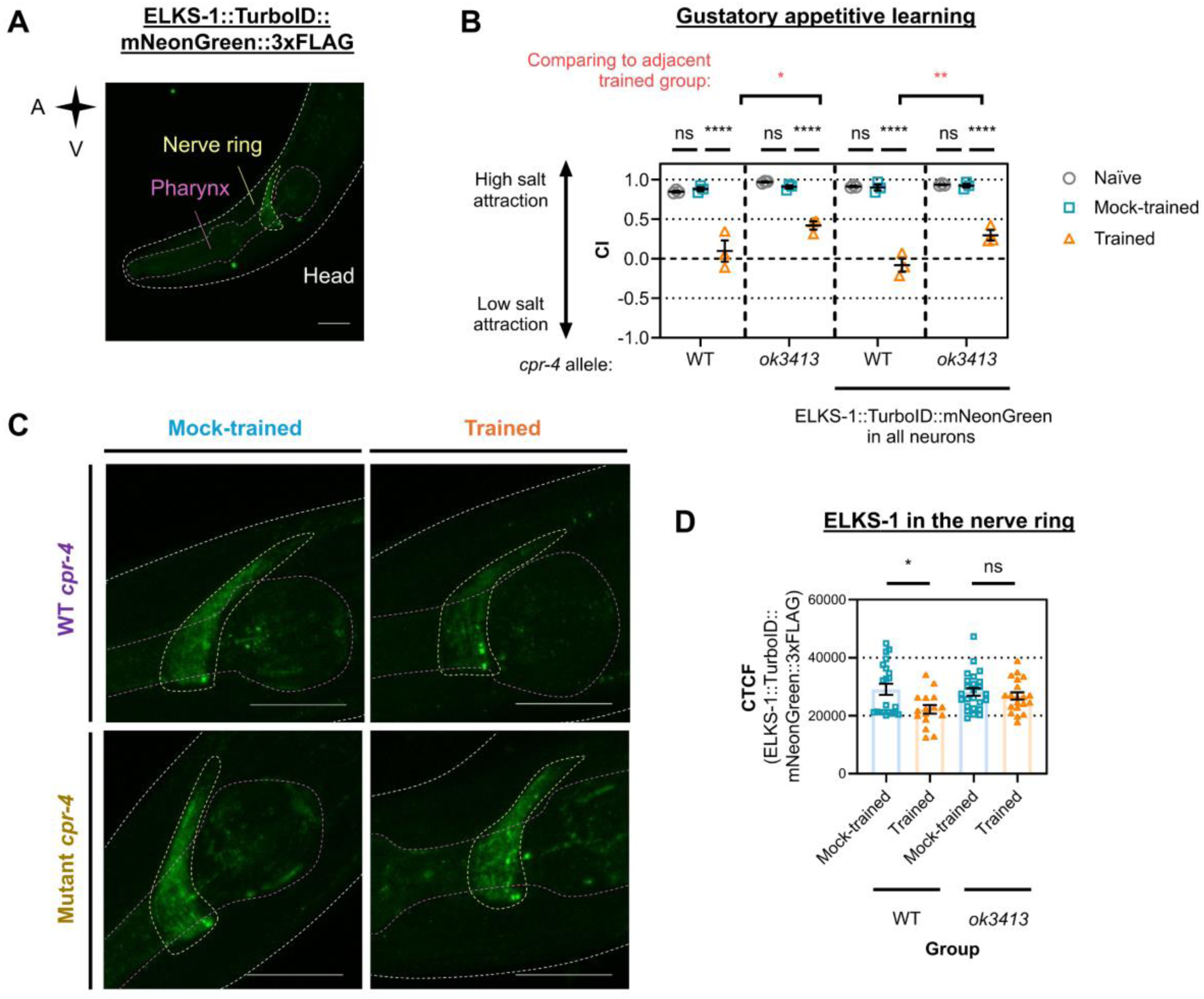
Learning-associated reduction of synaptic ELKS-1 requires CPR-4. **(A)** Representative image of a naïve *C. elegans* with mNeonGreen tagged ELKS-1 in a WT *cpr-4* background. ELKS-1 localises to the nerve ring (yellow outline), as previously reported [1]. Its body (white) and pharynx (purple) are labelled for anatomical reference. **(B)** Chemotaxis indices for animals mNeonGreen tagged ELKS-1, in a WT or *cpr-4(ok3413)* mutant background. Each data point represents one biological replicate (*n*). *n* = 3, technical replicates = 3 per *n*, 22-276 worms per technical replicate. Errors bars = mean ± SEM. Statistical analyses: Two-way ANOVA and Tukey’s multiple comparisons test. **(C)** Representative images of *C. elegans* after mock-training or gustatory appetitive learning. **(D)** Quantification of ELKS-1 in the nerve ring. Each data point is for a single animal (*n* = 16-25 per group). Statistical analyses: One-way ANOVA and Sidak’s multiple comparisons test. For all images, Anterior/A is left, ventral/V is down. Scale bar (white solid line) = 10 µm. p-values are indicated by: **** ≤ 0.0001, ** ≤ 0.01, * ≤ 0.05, ns = non-significant.

Next, we asked whether CPR-4 might directly interact with ELKS-1. We did not detect CPR-4 in co-immunoprecipitation experiments with ELKS-1, suggesting that the proteins do not stably interact under the conditions tested (**Fig. S5A & S5B**). We also investigated whether learning alters CPR-4 expression. We did not observe a training-dependent change in CPR-4 transcript levels, based on a 1.5-fold change threshold (**Fig. S5C & S5D**) [68]. Similarly, levels of secreted CPR-4, measured using CPR-4::mCherry fluorescence in coelomocytes, were unchanged (**Fig. S5E−S5G**). Together, these findings suggest that preexisting CPR-4 may influence ELKS-1-associated synaptic remodelling through an indirect mechanism, or through transient interactions not readily detected by co-immunoprecipitation, thereby contributing to learning-associated changes in synaptic protein composition (**Fig. 6**).

**Figure 6.**
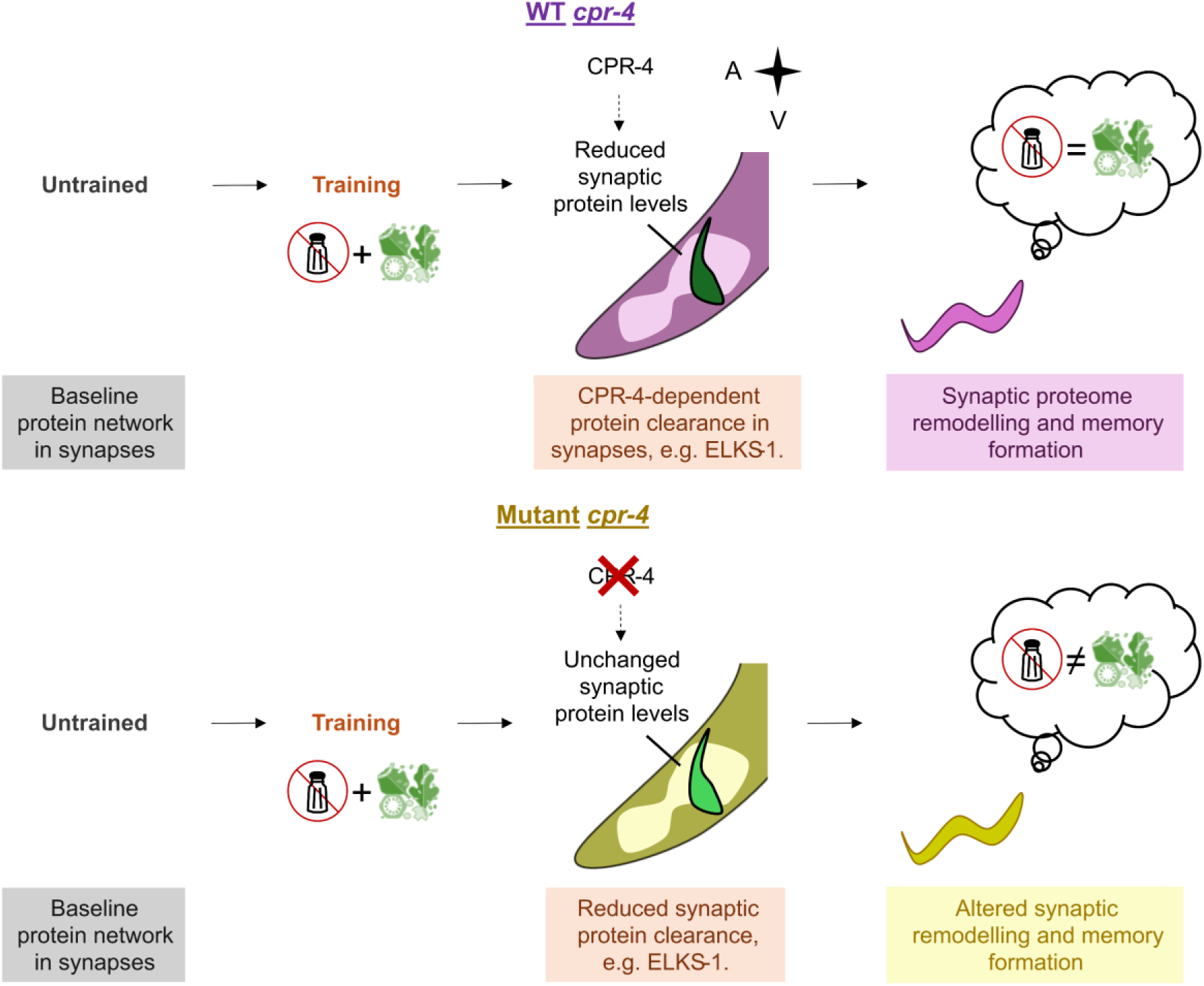
Proposed model for CPR-4-dependent synaptic remodelling during appetitive learning. Synapses contain a baseline network of proteins prior to training. We propose that appetitive learning induces remodelling of this protein network, in part through CPR-4 dependent protein clearance pathways. In wild-type (WT) animals (top), training is associated with reduced synaptic ELKS-1 levels, enrichment of protein turnover pathways in the synaptic proteome, and acquisition of a learning-associated behavioural and synaptic state. In *cpr-4* mutants (bottom), where memory formation is altered, learning-associated changes in synaptic protein composition are impaired, ELKS-1 levels remain high, and trained synapses retain features of the pre-learning state. Together, our findings support a model in which CPR-4 contributes to synaptic remodelling and memory formation through regulation of synaptic protein networks.

## Discussion

There is increasing evidence that synapses are remodelled at the protein level to encode memory. Using TurboID proximity labelling in synapses during gustatory memory formation, we found that trained synapses contained a protein network (∼300 proteins) that differed substantially from mock-trained controls. This included multiple proteins not previously studied in the context of learning, including CPR-4/CTSB. Through behavioural tests with single gene knockouts, we confirmed that CPR-4 is a novel regulator of learning. Further proteomic comparisons and targeted imaging experiments highlighted the role of CPR-4 in influencing the composition of synaptic proteins, shifting *C. elegans* neurons from a naïve to a trained proteome state. We propose a novel mechanism in which CPR-4 drives synaptic remodelling in appetitive learning, potentially through its predicted lysosomal protease activity.

To target *C. elegans* synapses during training, we employed TurboID fused to presynaptic scaffolding protein ELKS-1 in all neurons. Compared with non-tagged TurboID, which labels proteins throughout neurons [1, 23], targeting TurboID to synapses improved the enrichment and detection of biotinylated synaptic proteins (**Fig. 1C**). We initially intended to exclude all 123 proteins also detected by whole neuron TurboID during training from the synaptic learning proteome, as the enzyme traditionally serves as a ‘background’ control for proximity labelling [1, 24]. However, many overlapping proteins corresponded to known presynaptically localised mRNAs in the worm (i.e. ∼24%, [30]), suggesting that these are bona fide synaptic proteins or proteins that shuttle between synaptic and somatic components.

Many studies indicate that learning-associated synaptic plasticity is accompanied by extensive remodelling of synaptic protein composition. These changes occur through several complementary processes, including rearrangement of the actin cytoskeleton, regulation of neurotransmitter receptor abundance at synaptic membranes, and modification of synaptic protein composition through protein trafficking, translation, and degradation [19, 69–73]. Increasing evidence further suggests that protein turnover pathways contribute directly to learning-induced synaptic plasticity. In both flies and mice, learning is accompanied by increased autophagic flux and lysosomal activity, while lysosomal function is required for memory formation in *Drosophila* [74–76]. Consistent with this emerging view, our synaptic learning proteome was enriched for proteins and pathways associated with membrane trafficking, protein turnover, autophagy, and lysosomal function. Although we did not directly measure autophagic or lysosomal activity, the identification of CPR-4 together with multiple autophagy-associated proteins suggests that regulated protein turnover is a conserved feature of memory-associated synaptic plasticity. Importantly, while changes in synaptic protein composition with learning have been investigated in other systems, they have remained largely unexplored in *C. elegans*. By combining spatiotemporally resolved proteomics with behavioural genetics, we were able to characterise a learning-associated synaptic proteome and identify CPR-4 as a novel regulator of synaptic remodelling during learning.

One of the most striking findings of this study emerged from the similarity analyses of learning-associated synaptic proteomes. At both the protein and pathway levels, trained *cpr-4* mutants more closely resembled WT mock-trained than WT trained animals, suggesting that loss of CPR-4 prevents acquisition of the learning-associated synaptic state normally induced by training (**Fig. 4**). This effect was not simply due to a failure of learning, as WT and *cpr-4* mock-trained proteomes also differed, indicating that CPR-4 influences basal synaptic protein composition in addition to its role during learning. Collectively, these observations support a model in which CPR-4 contributes to transitions between synaptic proteome states. Pathway analyses further identified transcriptional regulation, protein turnover, and protein trafficking as candidate processes associated with CPR-4-dependent remodelling. Although these analyses are exploratory and do not necessarily indicate causal relationships, they suggest that CPR-4 may influence multiple aspects of synaptic plasticity at the protein level. Interestingly, studies of the mammalian orthologue CTSB have similarly implicated the protein in transcriptional regulation and ubiquitin-dependent protein turnover [77, 78], altogether suggesting some components of CTSB-family biology are evolutionarily conserved. As the processes through which CTSB influences learning in higher organisms remain unclear, our work provides a foundation for further mechanistic studies.

The learning-associated reduction in ELKS-1 abundance provides an example of a specific synaptic protein whose regulation depends on CPR-4. Because ELKS-1 is a core presynaptic scaffolding protein [25], changes in its abundance provide a useful readout of learning-associated synaptic remodelling. Trained WT animals exhibited reduced ELKS-1 fluorescence relative to mock-trained controls, and this reduction was absent in trained *cpr-4* mutants. These findings suggest that removal or redistribution of specific active zone proteins may contribute to learning-associated synaptic remodelling. Although we did not detect a stable interaction between CPR-4 and ELKS-1, transient interactions or indirect regulatory mechanisms remain possible. Together, these findings support our proteomic evidence that CPR-4 facilitates learning-associated changes in synaptic protein composition (**Fig. 5**), with ELKS-1 providing a potential molecular example of the proteome-state transitions identified in **Fig. 4**.

Because TurboID was targeted to ELKS-1, learning-associated changes in bait abundance must also be considered when interpreting the proteomic data. If reduced ELKS-1 abundance alone accounted for the proteomic differences observed between conditions, we might anticipate similar pathway compositions across proteomes, with fewer proteins detected in trained WT animals where ELKS-1 levels are lower. Instead, we find that learning alters both protein content and pathway composition in both WT and *cpr-4* mutant synapses (**Figs. 1C & 4C, Tables S7−S9**), indicating that the observed proteomic differences reflect bona fide changes in synaptic protein composition rather than differences in ELKS-1 abundance.

Another interesting finding was that CPR-4 exerted opposing effects on appetitive and aversive learning (**Fig. 3**). While CPR-4 promoted appetitive learning, it appeared to limit aversive learning, as *cpr-4* mutants displayed a stronger aversive learning response than WT animals. We did not investigate the molecular basis of this valence-dependent effect, and it remains unclear whether CPR-4 influences appetitive and aversive learning through a shared mechanism. Consequently, our proposed model is restricted to appetitive learning, where proteomic and imaging data indicate a role for CPR-4 in learning-associated synaptic remodelling. The aversive learning phenotype nevertheless raises the intriguing possibility that CPR-4 participates in additional signalling pathways that differentially regulate behavioural responses to positive and negative experiences. Interestingly, the mammalian orthologue, CTSB, may also exert context-dependent effects on cognition. For example, CTSB overexpression in skeletal muscle impaired spatial learning in mice [77], whereas exercise-induced increases in circulating CTSB were also positively correlated with memory performance in several human studies [79–81], but not universally (reviewed in [82]). Thus, neither CPR-4 nor CTSB appears to exert uniformly positive or negative effects on learning across all contexts. Notably, CPR-4 has previously been studied primarily as a stress-response factor induced by UV exposure or overcrowding [3, 60]. Its identification through learning-associated proximity proteomics therefore highlights the value of unbiased discovery approaches for uncovering previously unrecognised regulators of behaviour.

Several questions remain to be addressed. First, our proteomic strategy was qualitative and therefore identifies proteins associated with specific learning states rather than directly measuring abundance changes. Second, the proteins that are directly processed or regulated by CPR-4 remain unknown. Third, although our data support a role for CPR-4 in learning-associated synaptic remodelling and protein turnover, future studies measuring autophagic flux, lysosomal activity, and proteolytic function during learning will be important to establish the underlying mechanism. Finally, because ELKS-1 served as the TurboID bait and decreases during learning, future experiments using alternative synaptic baits will help determine the extent to which the identified proteome changes generalise across synaptic compartments. Despite these limitations, both protein- and pathway-level analyses support the conclusion that learning is accompanied by substantial remodelling of the synaptic proteome and that CPR-4 contributes to this process.

In summary, we identify CPR-4 as a regulator of synaptic proteome remodelling in learning for *C. elegans*. Our findings support a model in which CPR-4 promotes the transition between synaptic proteome states during appetitive learning, enabling animals to acquire a trained synaptic protein composition. By combining spatiotemporally resolved proximity proteomics, behavioural genetics, and functional validation, we generated the first learning-associated synaptic proteome for *C. elegans* and identified CPR-4 from hundreds of candidate proteins as a previously unrecognised regulator of learning. Given the high degree of conservation between *C. elegans* and humans, this work not only provides a framework for investigating how dynamic changes in synaptic protein networks contribute to memory formation, but also provides a valuable resource for uncovering conserved mechanisms for memory function.

## Methods

### C. elegans strain maintenance

Nematode maintenance was performed as in [23]. Gravid hermaphrodite animals at the first day of their adulthood were used for all experiments. They were grown in standard conditions on nematode growth medium agar at 22°C [83]. Proximity labelling was achieved with worms fed biotin-auxotrophic *E. coli* strain *MG1655 BioB::Kan* for two generations [20]. This was done to deplete biotin levels prior to the addition of biotin during training, to ensure strong temporal control of TurboID activity. *C. elegans* were otherwise given biotin-producing *E. coli* strain OP50 as their food source during their development. **Table S1** provides strain details for all *C. elegans* strains used in this study.

### Proximity labelling

The biotin treatment strategy used here was detailed in [23] and based on [1]. Biotin exposure was restricted to the 6 hr-time window that animals were mock-trained or trained by appetitive gustatory learning (defined below, see **Fig. 1A**). *C. elegans* were fed *E. coli* MG1655 in modified Luria Broth (LB; 25 mM NaCl, 5 mM K_3_PO_4_ (pH 6.0), 1 mM CaCl_2_, 1 mM MgSO_4_, 1.0% (w/v) Bacto^TM^ Tryptone, 0.5% (w/v) yeast extract, 0.05 mg/mL Kanamycin) for mock-training and training protocols. Bacteria were supplemented with 1 mM of exogenous biotin as its final concentration for proximity labelling, from a 1:100 dilution (originally in 250 mM KOH, 5 mM K_3_PO_4_ (pH 6.0), 1 mM CaCl_2_, 1 mM MgSO_4_). ≥3,000 animals per experimental group and biological replicate underwent this treatment before downstream proteomics.

### Protein extraction, protein quantification, and western blotting

Animals were washed twice with ‘worm washing buffer’ (WWB; 50 mM NaCl, 5 mM K_3_PO_4_ (pH 6.0), 1 mM CaCl_2_, 1 mM MgSO_4_) after mock-training/training. They were then sedimented by gravity while on ice to form a packed pellet, for storage in a –80°C freezer [23]. ≥3,000 animals constituted ∼300 µL of packed *C. elegans* in total.

Protein extraction methods were adapted from [84] and [1]. Each pellet was sonicated (10× 4 sec, 2 sec ‘on’ and 3 sec ‘off’) in 200 μL of Radioimmunoprecipitation assay (RIPA) buffer (2 M urea, 150 mM NaCl, 50 mM Tris-Cl (pH 8.0), 5 mM EDTA, 10 mM NaF, 2 mM Na_3_VO_4_, 1 mM NaPP, 1% (v/v) Nonidet-P40, 1% (w/v) SDS, 0.5% (w/v) sodium deoxycholate, 0.1% (w/v) β-glycerophosphate, 1×cOmplete Mini Protease Inhibitor (Merck)) [1, 85]. This was done with the Q125 Sonicator (Q Sonica) in a cold room (∼2–8°C), with a >20 sec recovery on ice in-between sonications to minimise sample degradation. All samples were vortexed for ∼5 sec at room temperature to further lyse *C. elegans*, and then centrifuged (14,000 rcf, 4°C, 10 min) to separate carcasses/debris from supernatant containing total protein (i.e. from whole worm bodies). Total protein samples were assessed by bicinchoninic acid assay as per manufacturer’s instructions (Thermo-Fisher Scientific, #23225).

Standard protocols were utilised for SDS-PAGE, semi-dry protein transfer, and western blotting [86–88]. Proteins of interest were probed with the antibodies/conjugates listed in **Table 1** and visualised by chemiluminescence using Clarity Western ECL Substrate (Bio-Rad, #1705060) according to manufacturer’s instructions.

**Table 1.** Probes used to visualise proteins of interest in western blot experiments. All antibodies and conjugates were prepared in 5% (w/v) BSA in TBS-T.

| Probe name | Dilution | Manufacturer | Catalogue number |
| --- | --- | --- | --- |
| Goat anti-rabbit HRP | 1:10,000 | ThermoFisher Scientific | #31460 |
| Rabbit anti-GFP | 1:1,000 | Cell Signalling Technology | #2956 |
| Rabbit anti-mCherry | 1:1,000 | Cell Signalling Technology | # 43590 |
| SA-HRP | 1:5,000 | Cell Signalling Technology | #3999 |

### Preparation of peptide samples for LC−MS/MS runs

Peptide samples were generated using protocols from [23, 24, 85, 89]. This was initially done with ∼0.6–1.0 mg of desalted total protein per experimental group and biological replicate in our discovery experiment in **Figure 1**. This starting material was increased to 2 mg for **Figure 4**, to improve biotinylation signal from synaptic TurboID. Total protein was desalted with 7 kDa molecular weight cut-off desalting spin columns (ThermoFisher Scientific, #89883) by buffer exchange with TBS-P (150 mM NaCl, 10 mM Tris (pH 7.4), 1×cOmplete Mini Protease Inhibitor) [23, 24]. We then performed two pull-downs sequentially with desalted total protein [24].

First, each sample (0.7 mL total volume) was gently agitated for 1 hr in a cold room (∼2–8°C) with nickel-nitrilotriacetic acid (Ni-NTA) agarose resin equilibrated with TBS-P (10:4 protein-to-bead ratio). This was done to selectively enrich proteins endogenously biotinylated in the worm using transgenic poly-histidine tags. The resin was spun down briefly with a mini-centrifuge to isolate supernatant containing total protein that did not bind to the resin minus poly-histidine tagged proteins, for LC−MS/MS. The remaining bound protein was further processed as described below.

Poly-histidine tagged proteins were washed twice with a cleansing buffer (150 mM NaCl, 50 mM Tris (pH 7.4), 1 mM imidazole, 0.1% (w/v) SDS), incubated in cleansing buffer supplemented with 500 mM imidazole at room temperature for 10 min, and then spun down by centrifugation (10,000 rcf, 5 min, 22°C). This was done to isolate supernatant containing eluted endogenously biotinylated proteins with poly-histidine tags. These proteins then underwent a second pull-down similar to that done using Ni-NTA, to concentrate proteins into a smaller volume that could be loaded into SDS-PAGE gels. This was done with Streptavidin Magnetic Beads (NEB, #S1420S) equilibrated with TBS-T by incubation for 18 hr (8.0:4.5 protein-to-bead ratio, 0.25 mL total volume per sample). Bound proteins underwent 3× TBS-T washes before they were denatured in 1× sample buffer (as described above). This protocol was provided by Dr Emmanuel Prikas (*pers. comm.,* and [89]).

Biotinylated proteins that did not bind to Ni-NTA resin were enriched for LC−MS/MS using Streptavidin Magnetic Beads. Magnetic beads were processed as in [23], with protocols originally made by [85] and [89]. They were washed with the following solutions (number of washes in brackets): TBST (×3), 1 M KCl (×1), 0.1 M Na_2_CO_3_ (×1), and PBS (×5; ThermoFisher Scientific, #10010023). Each sample was then split 1:9 for western blots (as above) and LC−MS/MS, respectively.

Enriched proteins for LC−MS/MS were shaken vigorously to promote reduction with 5 mM dithiothreitol (800 rpm, 55°C, 1 hr) and then alkylation with 0.01 M iodoacetamide (800 rpm, 55°C, 20 min in the dark) (in 2 M urea, 7 mM NH_4_HCO_3_, 0.1% (w/v) Protease-Max Surfactant (Promega, #V2071)). Afterward, proteins underwent on-bead digestion with 0.01 µg/µL of Sequencing Grade Modified Trypsin (Promega #V5111) (in 50 mM NH_4_HCO_3_ and 0.015% (w/v) Protease-Max Surfactant (Promega, #V2071)) via an 18 hr-incubation set to 800 rpm and 37°C. Trypsin was quenched as described previously with trifluoroacetic acid [23, 89], resulting in two peptide sample types for each experimental group and biological replicate: (1) biotinylated peptides eluted from beads as the ‘bound’ fraction, and (2) unbiotinylated peptides that naturally fell away from the beads during the trypsin digest as the ‘unbound’ fraction.

Peptides were desalted with tC18 cartridges (Waters, #WAT036810), vacuum-dried (room temperature, ∼3 hr), and then reconstituted in 0.2% (v/v) heptafluorobutyric acid in 1.0% (v/v) formic acid for LC−MS/MS [23, 89].

### LC−MS/MS runs

We used the ThermoFisher Scientific Orbitrap Exploris mass spectrometer to generate raw m/z peak data (*n* = 3 in **Fig. 1**; *n* = 2 in **Fig. 4**) as in [89]. m/z data was converted into protein identities using the MASCOT search engine (Matrix Science) and a *C. elegans* Swiss-Prot database [downloaded 19/01/2024]. We used the same settings as in our previous work to perform an MS/MS ion search with this software: ‘semi-trypsin’ enzyme, ‘monoisotopic’ mass values, ‘unrestricted’ protein mass, ‘±5 ppm’ peptide tolerance, ‘±0.05 Da’ fragment mass tolerance, and ‘3’ maximum missed cleavages. We selected Biotin (K), Carbamidomethyl (C), Oxidation (M), Phospho (ST), and Phospho (Y) as variable modifications [23]. We used the online tool STRING (v12) [accessed 05/07/2024] to convert protein IDs to the gene names shown in **Tables S2 & S6** [90]. MASCOT search results for this study are provided via the Dryad platform (see *Data Availability*).

### Analyses from LC−MS/MS data

#### Proteome list determination

Protein identities were compared using a qualitative approach [23]. This involved four consecutive steps for our discovery experiment (**Fig. 1**) where, for three initial steps, groups within the same biological replicate were compared together, and then we subsequently combined data across biological replicates:

1. **Combining unbound and bound peptide sample data for each biological replicate:** Protein identities determined from each peptide sample type for the same experimental group and biological replicate were combined. This generated the initial protein identity lists that we further processed as described below.
2. **Removing non-specific background from each biological replicate**: We compared no TurboID, whole neuron TurboID, and synaptic TurboID groups. To do this, we subtracted proteins from ‘whole neuron TurboID, mock-trained’ and ‘synaptic TurboID, mock-trained’ lists when they were detected in ‘no TurboID, mock-trained’. This was then repeated with the corresponding trained groups.
3. **Determining proteins unique to trained animals for each biological replicate**: Mock-trained proteins were deducted from trained protein lists, and vice versa.
4. **Combining biological replicates to identify whole neuron vs synaptic proteomes**: Finally, protein lists from each biological replicate were combined for TurboID groups, to output lists shown in **Table S2**.

A similar approach was used for LC−MS/MS experiments shown in **Figure 4**. These lists were then combined for all biological replicates (**Table S6**).

Venn diagrams were generated to compare proteome datasets using an online tool (https://bioinformatics.psb.ugent.be/webtools/Venn/). The overlap between proteome datasets in each Venn diagram represents proteins detected in each experimental group in different biological replicates, when the group was used as a background control during subtractions.

#### Tissue expression assessment

Each proteome list was assessed for nervous system expression using the adult CeNGEN database [26, 27]. We used the ‘gene expression by cell type’ function to bulk search gene names corresponding to every protein in a proteome dataset (threshold = 2). This data was used to calculate the percentage of neuron expressed proteins in a dataset, as those expressed in at least one neuron class vs the total number of proteins in the same dataset. The software used to perform this *in silico* analysis is available online (https://cengen.shinyapps.io/adult/).

#### KEGG, Reactome, and Biological Process database analyses

ShinyGO was used to determine KEGG and Reactome terms (v0.85.1, from 2025) [accessed 26/06/2026], and for Biological Process (v0.86.0, released 28/08/2026) [accessed 31/08/2026] [28, 29, 40–42]. Searches were performed with the ENSEMBL dataset for *Caenorhabditis elegans* nematodes containing N2 genes (WBcel235) with ENSEMBL ID ‘celegans_gene_ensembl’. No background information was uploaded for these searches, meaning fold enrichment calculations were based on the total number of *C. elegans* proteins with the same term. We set ‘FDR cutoff’ to 0.05, ‘number of shown pathways’ to 20, ‘minimum pathway size’ to 2, and ‘maximum pathway size’ to 5,000. We set only the ‘remove redundancy’ and ‘abbreviate pathway’ features as ‘on’ for each search. Then, for each protein list corresponding to the four proteomes above, .csv files containing all pathway information were exported from the ‘enrichment’ tab on ShinyGO (see **Tables S4−S5 & S7−S9**). This data was visualised using R (v4.6.0) [91] with packages dplyr [92], ggplot2 [93], readr [94], tidytext [95], and tidyverse [96].

#### Jaccard chart generation

We converted protein identity and Reactome pathway information to a binary matrix, indicating those present in each proteome dataset as 1 vs absent as 0. This information was converted into a Jaccard chart using R (v4.6.0) [91] and packages pheatmap [97], RColorBrewer [98], readr [94], tidyverse [96], and vegan [99].

### Co-immunoprecipitation

We adapted the immunoprecipitation protocols from [100] and [101]. ≥3,000 animals were pelleted per experimental group (∼300 µL pellet volume), for sonication in 2.0× lysis buffer (100 mM KCl, 60 mM HEPES (pH 7.4), 10 mM NaF, 4 mM MgCl_2_, 2 mM DTT, 2 mM Na_3_VO_4_, 1 mM NaPP, 10% (v/v) glycerol, 0.5% (w/v) sodium deoxycholate, 0.1% (w/v) β-glycerophosphate, 0.1% (v/v) TritonX, 1×cOmplete Mini Protease Inhibitor). Protein G Magnetic Beads (NEB, #S1430S) were used to preclear 1 mg total protein per group via a 1 h-incubation on an active tube rotator in a cold room (∼2–8°C). Proteins that were non-specifically bound during this preclearing step serve as ‘preclear’ samples. Proteins that did not bind non-specifically to protein G during preclearing (i.e. in the supernatant) were used for co-immunoprecipitation. To do this, each sample was gently agitated overnight (∼16 h) at ∼2–8°C with protein G beads and 1 μg of monoclonal mouse GFP antibody simultaneously (Merck, #11814460001). Both preclearing and co-immunoprecipitation steps were done with a 50:1 protein-to-bead ratio with 1.0× lysis buffer-equilibrated beads. Proteins that did not bind to protein G in the presence of our GFP antibody were separated from the magnetic beads as ‘supernatant’ samples. The remaining immunoprecipitated (IP) proteins were washed thrice in co-IP washing buffer (100 mM KCl, 30 mM HEPES (pH 7.4), 2 mM MgCl_2_, 1 mM DTT, 10% (v/v) glycerol, 0.1% (v/v) TritonX).

### RNA extraction and qPCR to quantify cpr-4 transcripts

RNA was extracted from whole animal bodies with a previously established method [68]. ∼500 nematodes (∼50 µL total volume) were prepared for RNA extraction immediately after mock-training/training, by pelleting them for freezing at −80°C. They were then mechanically lysed in Monarch StabiLyse DNA/RNA Buffer with 5 mm stainless steel beads on a QIAGEN TissueLyser at 30 Hz for 4 min. RNA was extracted from this lysate using a Monarch® Spin RNA Isolation Kit (NEB, #T2110) using the manufacturer’s protocol.

We used the Luna® Universal One-Step RT-qPCR Kit (NEB, #E3005) as in manufacturer’s instructions to conduct qPCR runs. Transcript information was measured using the QIAGEN Rotor-Gene Q real-time PCR machine. See **Table 2** for primer details.

**Table 2.**
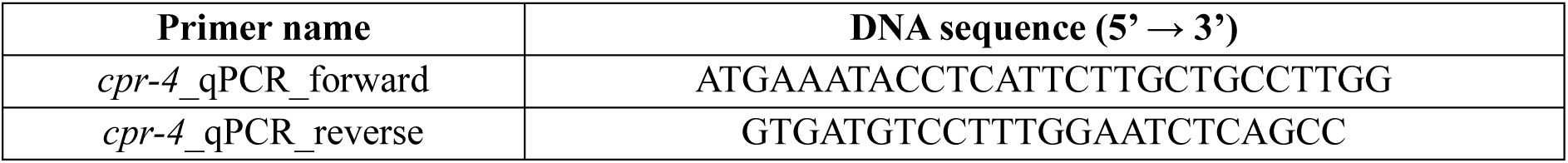
qPCR primers to measure *cpr-4* transcripts in mock-trained vs trained nematodes. We designed DNA primers to amplify a 143 bp product beginning from AUG in *cpr-4* mRNA corresponding to its start codon. They were designed using online tools ‘Multiple Primer Analyzer’ (ThermoFisher Scientific) and ‘NEB melting temperature calculator’ (v1.17.0).

### Gustatory appetitive learning

This behavioural test, referred to as ‘salt associative learning’ in [23], was originally adapted from [102]. We tested each genotype for their ability to learn by generating three experimental groups: naïve/untrained, mock-trained (50 mM NaCl + food), and trained (0 mM NaCl + food). Worms were collected from nematode growth medium plates using WWB and then washed twice with the same buffer. At this stage, animals were naïve and could be assessed for their behavioural response toward salt. *C. elegans* collected for training were washed a third time with no salt buffer (NSB; 5 mM K_3_PO_4_, 1 mM CaCl_2_, and MgSO_4_) to deplete NaCl levels. They were then transferred onto solid media containing 2% (w/v) agar, 5 mM K_3_PO_4_, 1 mM CaCl_2_, and MgSO_4_ in 90 mm diameter petri dishes. This agar was seeded with *E. coli* MG1655 in modified Luria Broth. Animals underwent a 6 hr-training period at 22°C in this environment. Mock-training was performed similarly but with WWB for the third wash and solid media supplemented with 50 mM NaCl for the 6 hr-incubation (i.e. the same concentration as during cultivation). WWB was used to collect mock-trained and trained animals. They were washed once with WWB before commencing chemotaxis assays.

### Gustatory aversive learning

This behavioural test, referred to as ‘salt aversive learning’ in [23] and [103], was originally adapted from [64].We prepared three experimental groups per genotype as in the ‘salt aversive learning’ section in [23]: naïve, mock-conditioned (0 mM NaCl + no food), and conditioned (50 mM NaCl + no food). Animals were collected and washed as for appetitive learning, but the third wash involved NSB for mock-conditioned and WWB for conditioned. Animals were agitated in liquid media within 1.5 mL tubes using an orbital shaker (175 rpm, room temperature, 3 hr) to facilitate mock-conditioning in NSB or conditioning in WWB. They were allowed to sediment by gravity for 1 min before use in chemotaxis assays.

### Salt chemotaxis assay

Animals were allowed to explore a salt concentration gradient (0-200 mM NaCl) in chemotaxis assay agar (2% (w/v) agar, 5 mM K_3_PO_4_, 1 mM CaCl_2_, and MgSO_4_) in 60 mm diameter petri dishes for 45 min at 22 °C [102]. Salt gradients were prepared with two 5 mm cubes of chemotaxis assay agar, where one cube was supplemented with 200 mM NaCl. To do this, two cubes were left overnight at 22 °C equidistant from each other on the agar surface of each plate [104]. Animals were paralysed at extreme ends of the salt gradient by 100 mM NaN_3_ (paralytic agent) during the 45 min-incubation. This allowed animals to be counted in defined regions to quantify their behaviour as a chemotaxis index:

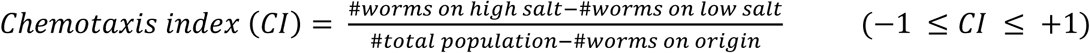

This protocol is described in detail in [23, 103].

### Imaging experiments

#### Microscopy

*C. elegans* undergoing mock-training/training by gustatory appetitive learning were transferred onto a freshly made 2% (w/v) agarose pad in a paralytic agent. Animals were paralysed in 100 mM NaN_3_ within 1 min (*n* = 3, **Fig. 5**) or 5 mM in tetramisole after 10 min (*n* = 1, **Fig. S5**) [105, 106].

Confocal imaging was performed using a Zeiss LSM 880 confocal microscope and 63× objective (NA: 1.4). Fluorescence settings were optimised with a control group and then kept consistent for all groups, i.e. mock-trained ELKS-1::TurboID::mNeonGreen nematodes (**Fig. 5**) or naïve CPR-4::mCherry worms (**Fig. S5G**). Z-stacks set to 1.00 μm steps (**Fig. 5**) or 0.75 μm steps (**Fig. S5G**) included one slice without fluorescence above and below each animal to ensure all fluorescent signal was captured. Images were converted to maximum intensity z-projections for quantification using the Fiji software (v2.9.0).

Coelomocyte imaging (**Figure S5F**) was performed with the Zyla sCMOS camera (Andor Technology), Nikon Eclipse Ti2 microscope, and imaging software micro-manager (v2.0.0). Snapshots were taken with a 0.034 sec capture speed and 100% intensity.

Fluorescence was quantified using ImageJ 1.53k software (Java, v1.8.0) based on [107]. To do this, the freehand selection tool was utilised to manually outline the nerve ring (**Fig. 5A**) or coelomocytes (**Fig. S8F**) in nematode images, and the ‘measure’ function was used to output integrated fluorescence density and area measurements. This was done for each individual image for regions outside of the head three times to calculate the average ‘mean grey value’ or fluorescence from background readings. Corrected total cell fluorescence (CTCF) values were calculated using the formula below:

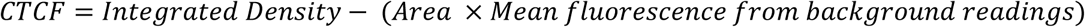

### Statistical analyses

#### Behavioural experiments

Five biological replicates were performed for each behavioural test in **Figure 3**. We did not blind genotype or condition information during chemotaxis assays. Technical replicates with individual animals less than 20, or evident bacterial contamination, were excluded from this study. All experiments were assessed by Two-way ANOVA and Tukey’s multiple comparisons test in line with previous studies [108–110].

#### Imaging

Single animals represent technical replicates for *n* = 3 in **Figure 5** and *n* = 1 in **Figure S8** (see figure legends for details). Experiments were blinded during CTCF quantification. For ELKS-1::TurboID::mNeonGreen, we excluded all images where the nerve ring was not orientated directly sideways, as the nerve ring is not symmetrical and therefore including all angles would confound results. Experimental groups were assessed by one-way ANOVA and Sidak’s multiple comparisons test for ELKS-1::TurboID::mNeonGreen. mCherry experiments included two experimental groups so they were compared with an unpaired two-tailed t-test using GraphPad Prism (v8.0).

## Data availability

*C. elegans* lines made by the Worm Neuroscience Lab (Flinders) for this publication are available upon request. The raw data from mass spectrometry and behavioural experiments in this work can be accessed using the Dryad database, which will be made available following publication.

## Acknowledgements

We thank colleagues in the Worm Neuroscience lab (Flinders) for important discussions, and acknowledge individuals who contributed to experiments in this study: Dr Radwan Ansaar for assisting in *C. elegans* maintenance, Pasquale Vitaro in preparing agar plates for behavioural experiments, and Khadija Dar for analysis of mCherry fluorescent data. We thank A/Prof Arne Ittner, Dr Emmanuel Prikas, and Dr Nicholas Eyre for scientific advice. We also acknowledge Flinders colleagues Prof Briony Forbes, Prof Kim Hemsley, A/Prof Mary-Louise Rogers, A/Prof Arne Ittner, and Dr Amy Wyatt for sharing reagents. We gratefully acknowledge Prof John E Cronan (University of Illinois, USA) for providing us with *E. coli* strain MG1655. We sincerely thank Prof Mario de Bono (Institute of Science and Technology, Austria), Prof Ding Xue (University of Colorado, USA), and the *Caenorhabditis* Genetics Centre supported by the National Institutes of Health (P40 OD010440) for supplying many *C. elegans* strains used in this study.

## Author contributions

Conceptualization, A.R. and Y.L.C.; Methodology, A.R. and Y.L.C.; Data curation, A.R., A.P., and Y.L.C.; Formal analysis, A.R., M.E.J., R.A.C., A.P., and Y.L.C.; Investigation, A.R., M.E.J., R.A.C., N.H., and A.P.; Writing—Original Draft, A.R. and Y.L.C.; Writing—Review & Editing, A.R., M.E.J., and Y.L.C.; Visualisation, A.R. and Y.L.C.; Funding acquisition, Y.L.C.; Project administration, Y.L.C; Supervision, Y.L.C.

## Ethics considerations

No ethics approval or welfare guidelines were required since *C. elegans* is a non-protected invertebrate species. *C. elegans* work was conducted under an approved exempt dealing authorised by the Flinders University Institutional Biosafety Committee.

---

**Figure S1.**
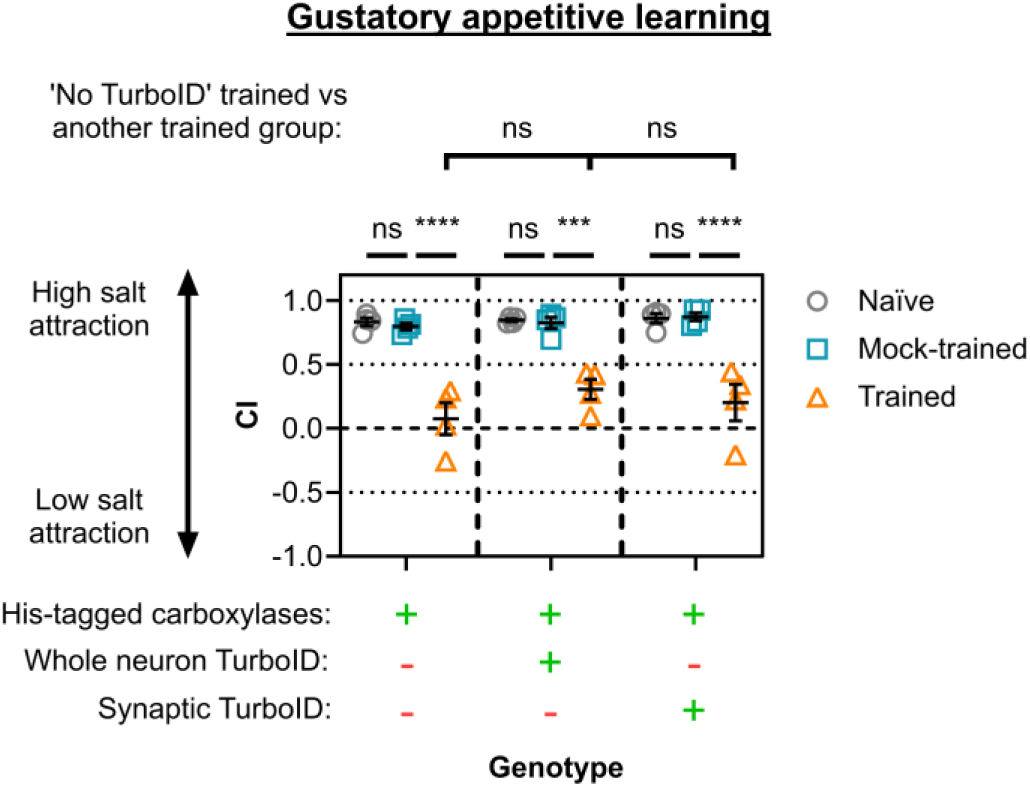
Proximity-labelling machinery does not impair gustatory learning in the *C. elegans*. Chemotaxis indices (CIs) for proximity labelling strains used in Figure 1. Salt chemotaxis behaviour was quantified using the CI (see *Methods*). Each data point represents one biological replicate (*n* = 4) each from three technical replicates (20-257 worms per technical replicate). Errors bars = mean ± SEM. Statistical analyses done with Two-way ANOVA and Tukey’s multiple comparisons test (**** ≤ 0.0001, *** ≤ 0.001, ** ≤ 0.01, * ≤ 0.05, ns = non-significant).

**Figure S2.**
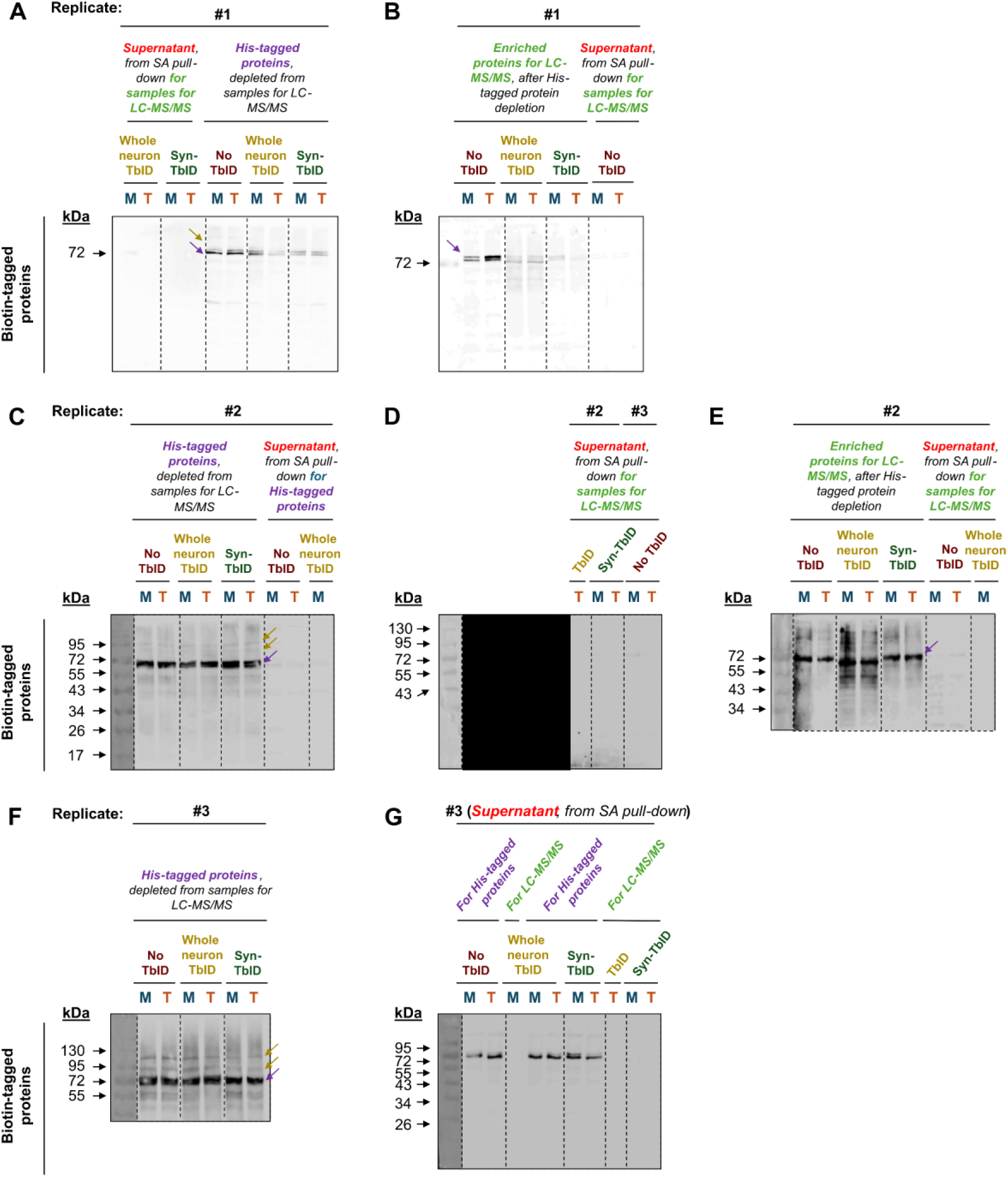
Double pull-down enrichment strategy for proximity-labelled proteins. **(A, C, F)** Western blots showing endogenous biotinylated and histidine-tagged carboxylases enriched during pull-down #1 using nickel-NTA. Bound proteins are shown in the **His-tagged protein** lanes (purple). **(B, E)** Western blots showing biotinylated proteins enriched during pull-down #2 using streptavidin. Bound proteins are shown in the **Enriched proteins for LC-MS/MS** lanes (green). **(A, B, D, E, G)** Western blots showing proteins remaining in the supernatant following nickel-NTA pull-down #1 and/or streptavidin pull-down #2. Supernatant fractions are shown in red. M = mock-trained, T = trained. Total protein underwent two pull-downs: (1) nickel-NTA affinity purification to remove histidine-tagged carboxylases and (2) streptavidin affinity purification to enrich biotinylated proteins. Yellow arrows indicate complete depletion of carboxylases, whereas purple arrows indicate partial depletion.

**Figure S3.**
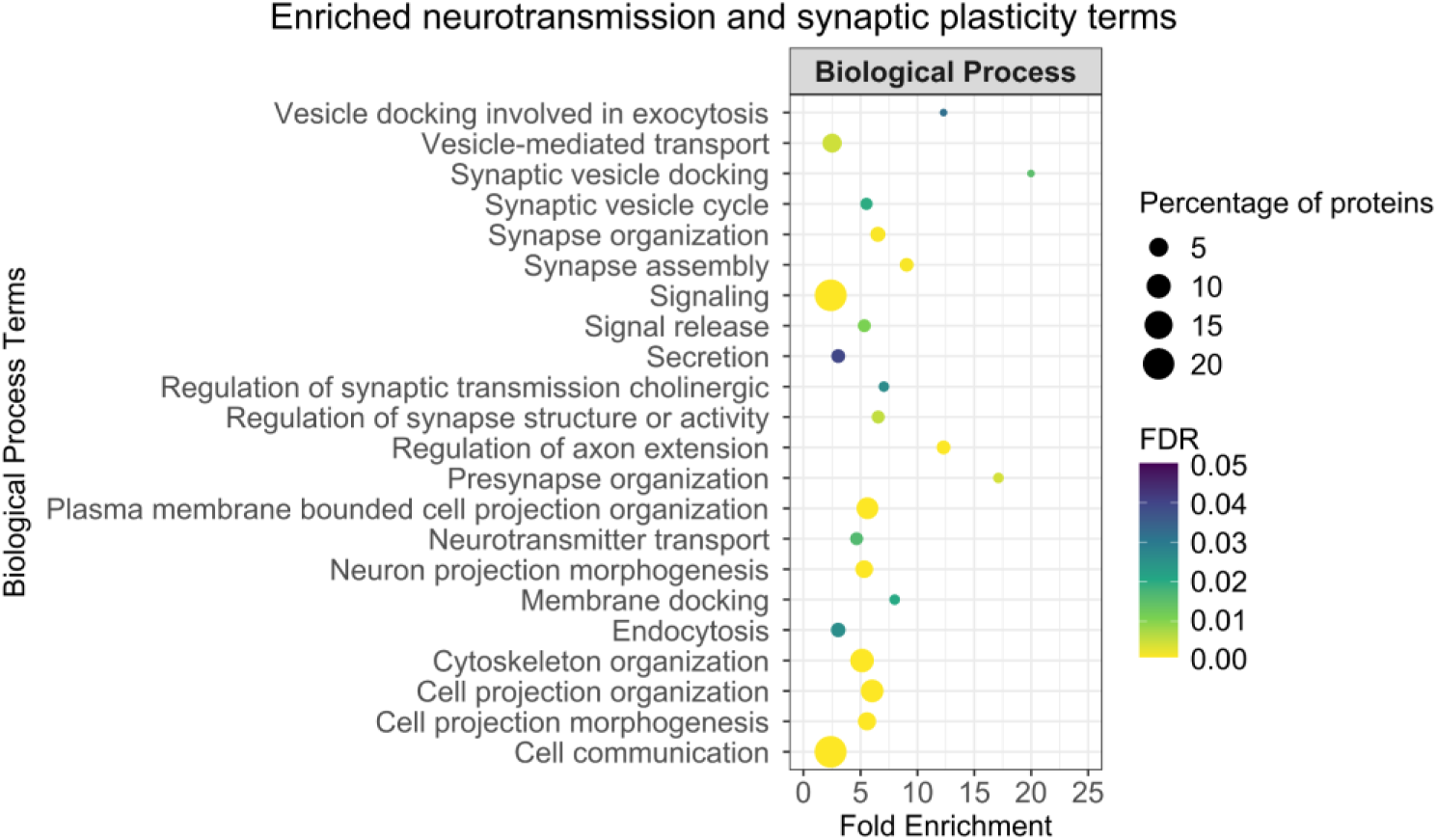
The synaptic learning proteome is enriched for neurotransmission and synaptic plasticity-related processes. Gene Ontology Biological Process enrichment analysis was performed using ShinyGO (v0.86) and visualised with R (v4.6.0). Terms related to neurotransmission regulation, protein trafficking to the synapse, and cytoskeletal reorganisation were selected for display. Node size represents the percentage of proteins from the synaptic learning proteome (319 proteins total) associated with each term. FDR ≤ 0.05.

**Figure S4.**
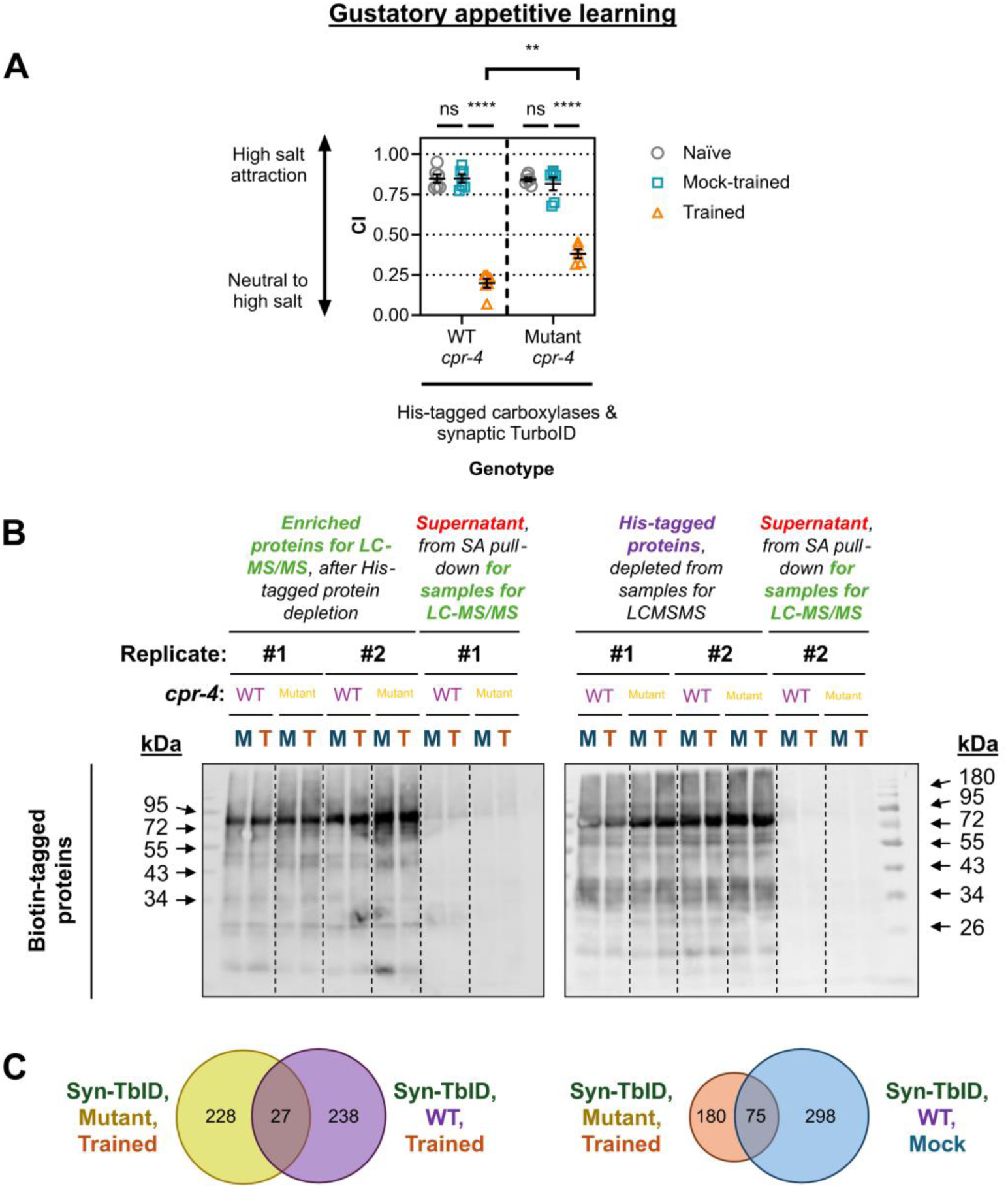
Synaptic TurboID successfully labels proteins in WT and *cpr-4* mutant animals. **(A)** Chemotaxis indices for strains expressing with His-tagged carboxylases and synaptic TurboID. Each datapoint represents a technical replicate (5-6 technical replicates in total per group). 2-3 technical replicates per *n* (*n* = 2) due to technical reasons, i.e. one technical replicate from mutant trained was excluded due to contamination of a chemotaxis assay plate. 92-458 worms per technical replicate. Errors bars = mean ± SEM. Statistical analyses: Two-way ANOVA and Tukey’s multiple comparisons test (**** ≤ 0.0001, ** ≤ 0.01, ns = non-significant). **(C)** Western blot analysis of biotinylated proteins following the double pull-down enrichment procedure (*n* = 2). Total protein underwent sequential enrichment (1) to deplete His-tagged proteins using Ni-NTA, and then (2) to enrich biotinylated with streptavidin in pull-down. Unbound proteins following streptavidin enrichment are referred to as ‘supernatant’. **(D)** Comparison of protein identities in synaptic proteomes datasets. Percentage overlap was calculated as the number of shared proteins divided by the total number of proteins represented in each comparison.

**Figure S5.**
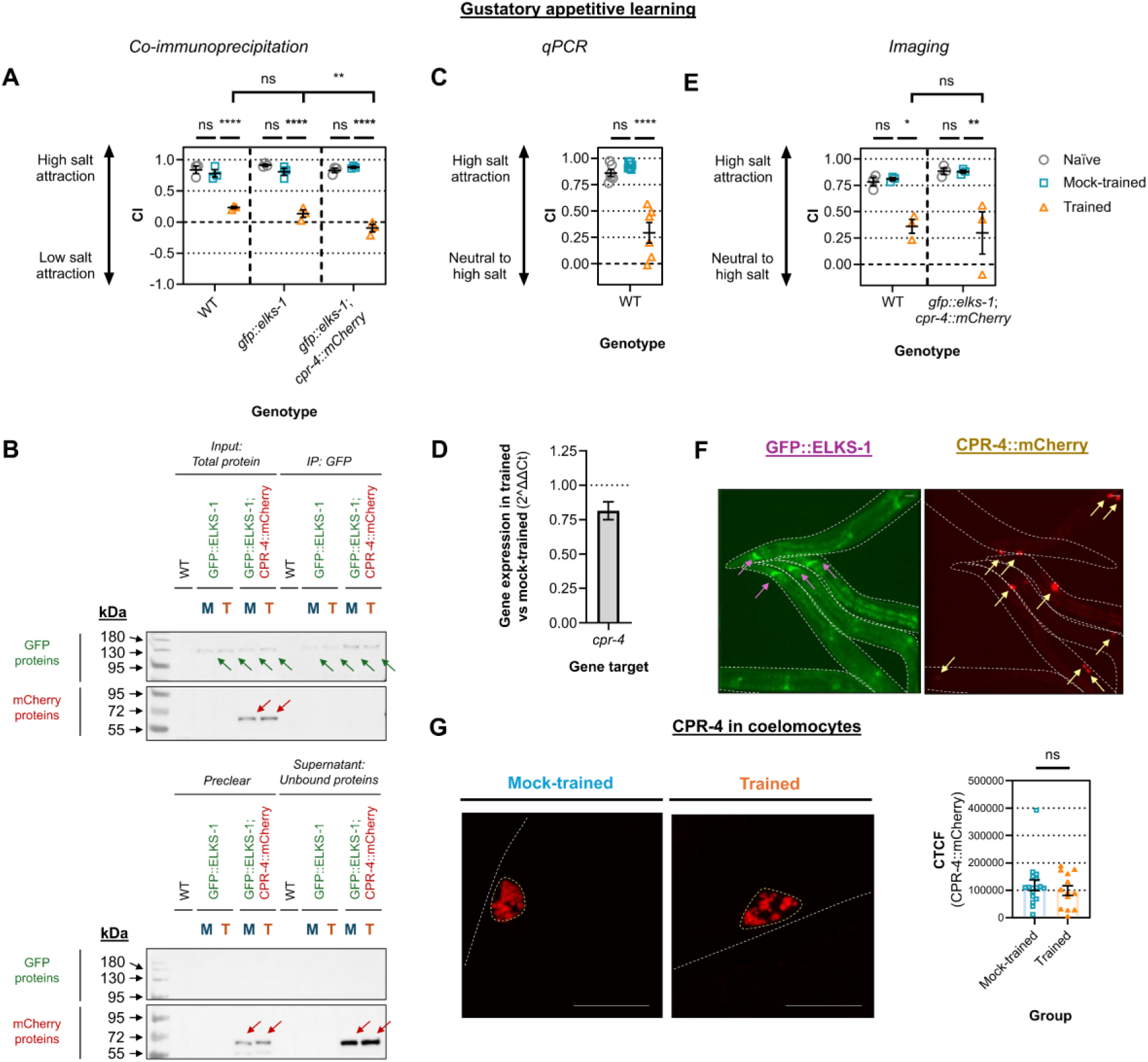
Control experiments examining CPR-4 expression and interaction with ELKS-1. **(A, C, E)** Chemotaxis indices for animals used in co-immunoprecipitation (*n* = 1, **A, B**), qPCR (*n* = 2, **C, D**), and imaging (*n* = 3, **E-G**) experiments. Technical replicates per *n* = 3, 43-266 worms per technical replicate. Each data point is one technical replicate **(A, C)** or one biological replicate **(E)**. **(B)** Co-immunoprecipitation analysis of ELKS-1 and CPR-4 in mock-trained (M) and trained (T) animals. Input lanes contain 40 µg of total protein. Preclear lanes show proteins that non-specifically bound to Protein G beads before the IP. IP lanes contain proteins enriched with a GFP antibody and Protein G beads. Supernatant lanes contain remaining unbound proteins after immunoprecipitation. Expected sizes: GFP::ELKS-1 = ∼130 kDa (green arrows), CPR-4::mCherry = ∼69 kDa (red arrows), calculated using ExPASy [16]. **(D)** qPCR analysis of *cpr-4* mRNA transcript abundance in WT animals. Fold change was calculated using the 2^ΔΔCt method, which represents the change in *cpr-4* gene expression in trained vs mock-trained controls. Errors bars = mean ± SEM. **(F, G)** Analysis of secreted CPR-4::mCherry in coelomocytes. **(F)** Representative images of naïve animals, with GFP::ELKS-1 in nerve rings (left, purple arrows) and CPR-4::mCherry in coelomocytes (right, yellow arrows). **(G)** shows maximum intensity z-projections of coelomocyte-localised CPR-4::mCherry in mock-trained vs trained *C. elegans* on the left. Scale bar = 10 µm. Quantification of mCherry signal is shown on the right as CTCF values. Errors bars = mean ± SEM. Data points represent individual coelomocytes (13-17 /group) from single animals (*n* = 9 /group) in one biological replicate.

